# Myosin turnover controls actomyosin contractile instability

**DOI:** 10.1101/2021.03.18.436017

**Authors:** Sathish Thiyagarajan, Shuyuan Wang, Ting Gang Chew, Junqi Huang, Mohan K. Balasubramanian, Ben O’Shaughnessy

## Abstract

Actomyosin contractile force is harnessed for diverse functions, from cell division to morphogenesis during development. However, actomyosin contractility is intrinsically unstable to self-reinforcing spatial variations that destroy actomyosin architecture if unopposed. The full instability was rarely observed, and how cells control the instability is not established. Here, we observed the instability run its full course in isolated cytokinetic contractile rings lacking component turnover. Myosin II aggregated hierarchically into aggregates of growing size and separation up to a maximum. Molecularly explicit simulations reproduced hierarchical aggregation that precipitated tension loss and ring fracture, and identified the maximum separation as the length of actin filaments mediating mechanical communication between aggregates. Late stage simulated aggregates had aster-like morphology with polarity sorted actin, similar to late stage actomyosin systems *in vitro*. Our results suggest myosin II turnover controls actomyosin contractile instability in normal cells, setting myosin aggregate size and intercepting catastrophic hierarchical aggregation and fracture.

## Introduction

Many fundamental single cell and tissue level processes rely on force production by actomyosin assemblies. During cytokinesis the actomyosin contractile ring develops tension that guides or drives furrow ingression and physical division of the cell^1, 2^. Tension gradients in the actomyosin cortex produce cortical flows that establish cell polarity^3^ and cortical actomyosin forces are harnessed for cell migration^4^. Contraction of supracellular actomyosin networks drives early tissue morphogenetic events such as gastrulation in *Drosophila*^5^ and *C. elegans*^6^ and neurulation in vertebrates^7^.

In these systems, actomyosin contractility finds use as a powerful and highly adaptable tool, in which contractile stress generated by constrained myosin II molecules that bind and exert force on oriented filamentous actin is harnessed for diverse functions. However, the mechanism has an intrinsic instability. A chance fluctuation in the density of myosin II and other components may enhance local contractile stress and draw in more actomyosin material at the expense of weaker neighboring regions, further amplifying the stress difference in a potentially runaway aggregation process (Fig. 1a). Unchecked, catastrophic fracture and tension loss may result.

**Figure 1.**
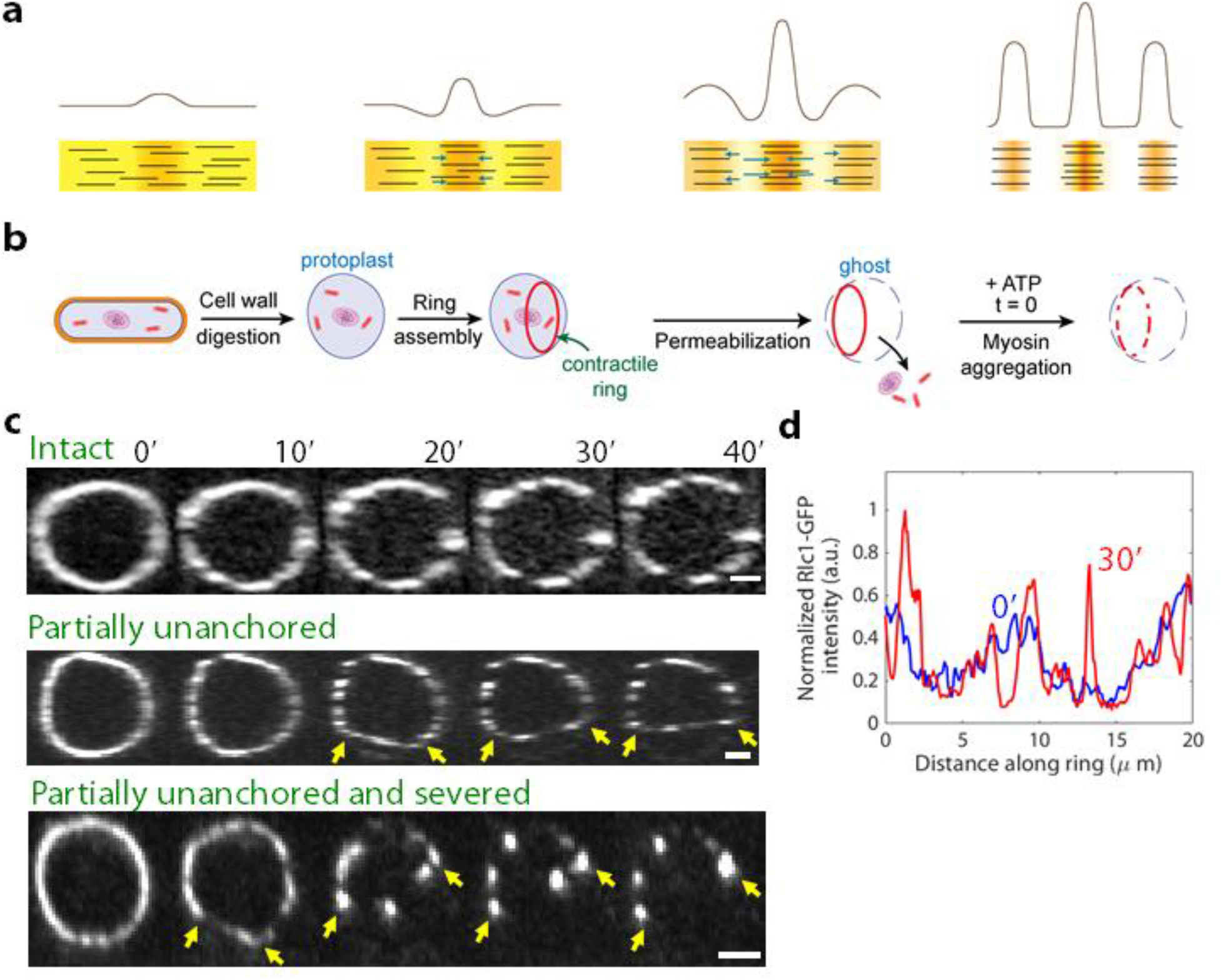
Aggregation of Myosin II in contractile rings of cell ghosts. (a) Actomyosin contractile instability, schematic. A fluctuation in the local density of contractile actomyosin material (left) generates contractile stress gradients and flows (arrows) that pull material inwards and amplify the fluctuation. Unopposed, the result is runaway aggregation and fracture (right). Density profiles are shown above schematics of the actomyosin material. (b) Preparation of cell ghosts. Following enzymatic digestion of cell walls of normal cells of the fission yeast *S. japonicus*, the plasma membrane is permeabilized to release cytoplasm. In cells which have assembled cytokinetic rings, the rings remain anchored to the membrane. On addition of ATP, myosin-II in rings aggregates progressively. (c) Time-lapse fluorescence micrographs of *S. japonicus* cell ghosts labelled with Rlc1-GFP following addition of 0.5 mM ATP. Imaging commenced at least 1-2 mins after ATP addition (see Methods). Contractile rings remain fully anchored to the membrane (top), or ring segments appear to detach from the membrane and shorten (middle), or segments detach, shorten and sever (bottom). Arrows indicate detached segments. We observed *n = 6, 5,* and *16* rings in each category, respectively. Scale bars, 2 μm. (d) Rlc1-GFP intensity along the ring at the indicated times for the intact ring of (b). Intensity normalized by the total intensity around the ring at each time.

How the cytokinetic contractile ring and other actomyosin machineries deal with this threat is not established. Some examples of the instability have been documented. In cells, actomyosin stress fibers spontaneously sever or undergo repair in regions of high strain^8, 9^, and *in vitro* myosin II spontaneously aggregated into puncta in reconstituted actomyosin bundles^10^. Theoretical evidence for contractile instability is abundant, as in molecularly explicit^11^ and continuum^12, 13^ mathematical models of the fission yeast contractile ring, and continuum models of the animal cell cortex^14, 15^.

Observing actomyosin contractile instability run its full course in cells has proved challenging, presumably because mechanisms intervene before its full maturation. Nevertheless, numerous observations are suggestive of incipient instability. Large fluctuations and punctateness characterize myosin II distributions in fission yeast and animal cell contractile rings^16, 17^ and animal cell cortices^18, 19^. Punctate cortical myosin in some cases has pulsatile time dependence over seconds to minutes^20^, which may be associated with pulsatile dynamics of the GTPase RhoA that promotes myosin II activity and minifilament assembly ^3, 15, 21^. Many morphogenetic events feature pulsed actomyosin epithelial contractions with punctate myosin II distributions^5, 22^.

Several candidate mechanisms that could control actomyosin contractile instability have been proposed. One possibility is turnover of myosin and actin^23, 24^, which presumably tends to homogenize spatial variations^11^. Another suggestion is myosin II diffusivity, which could serve this purpose by smoothing small scale density variations characteristic of the instability^14, 25^. Another proposal, suggested for *C. elegans* zygotes, is that the instabilities are controlled by pulsatile RhoA dynamics^15^.

Here we examined the role of component turnover as a possible regulator of actomyosin contractile instability by studying cytokinetic contractile rings in cell ghosts under conditions where turnover is absent^26, 27^. We find runaway aggregation of myosin II, in which the amount of myosin per aggregate and the separation between aggregates increases with time, up to a certain maximum separation. The aggregation is hierarchical, with repeated rounds of aggregation yielding aggregates with yet more myosin. In molecularly explicit simulations the maximum separation was identified as the length of actin filaments which mediate mechanical communication between aggregates, and small scale instability was controlled not by myosin diffusivity but by local excluded volume and actin filament polarity sorting effects. Our results suggest that in normal cells myosin II turnover controls contractile instability in actomyosin assemblies, setting the size of myosin II aggregates and preventing catastrophic hierarchical aggregation and fracture.

## Results

### Component turnover is absent in contractile rings of cell ghosts of the fission yeast *S. japonicus*

To investigate whether component turnover plays a role in maintaining organization in the cytokinetic contractile ring we used cell ghosts of the fission yeast *S.japonicus*. Following preparation of protoplasts by enzymatic digestion of cell walls, the plasma membranes of mitotic cells were permeabilized to release the cytoplasm and organelles, leaving isolated membrane-bound cytokinetic rings in cell ghosts^26, 27^ (Fig. 1b and Methods). Since cytoplasm is absent there is no component association, while dissociation rates are much reduced relative to normal cells^27^. Thus, cell ghosts allow the organization of isolated contractile rings to be studied in the absence of component turnover.

### In contractile rings without turnover, myosin II aggregates into puncta with increasing amounts of myosin

We imaged cells expressing myosin-II light chain Rlc1-GFP. Following membrane permeabilization the myosin II distribution was mildly punctate. On incubation with ATP, the distribution became progressively more punctate over ~30 min (Fig. 1c), consistent with our earlier study^27^. As the punctateness increased, contractile rings suffered three distinct fates (Fig. 1c). Some rings appeared to remain robustly anchored to the membrane, with myosin II aggregates located around the contour defined by the initial ring. In other rings, segments appeared to detach from the membrane and straighten, presumably due to compromised anchoring to the membrane^28^. In a third category, segments detached from the membrane and severed, following which myosin puncta in the severed segment merged with the apparently anchored segment.

For each ring we identified the longest segment attached to the membrane that neither severed nor underwent large length changes, and we measured the Rlc1-GFP intensity along the segment. Following ATP addition, peaks of myosin fluorescence intensity appeared which became increasingly prominent with time (Fig. 1d). The relative peak amplitudes increased with time while intensity decreased in regions neighboring the peaks. The peak intensities significantly exceeded the initial intensity, indicating that the puncta were at least in part due to movement of myosin around the ring.

Thus, myosin progressively merges into aggregates of growing size in contractile rings lacking component turnover. This behavior would be expected of unopposed actomyosin contractile instability, suggesting that turnover may control the instability in normal cells.

### Myosin II aggregates hierarchically in the absence of turnover

Kymographs of myosin fluorescence intensity profiles around contractile rings showed that, in regions where the myosin density was initially above the mean, the density increased in time and the region of higher density became smaller (Fig. 2a). In regions with an initial deficit the density decreased and the region expanded. Thus, excess density fluctuations grew and sharpened into distinct aggregates, while deficit density fluctuations diminished and became empty regions separating aggregates.

**Figure 2.**
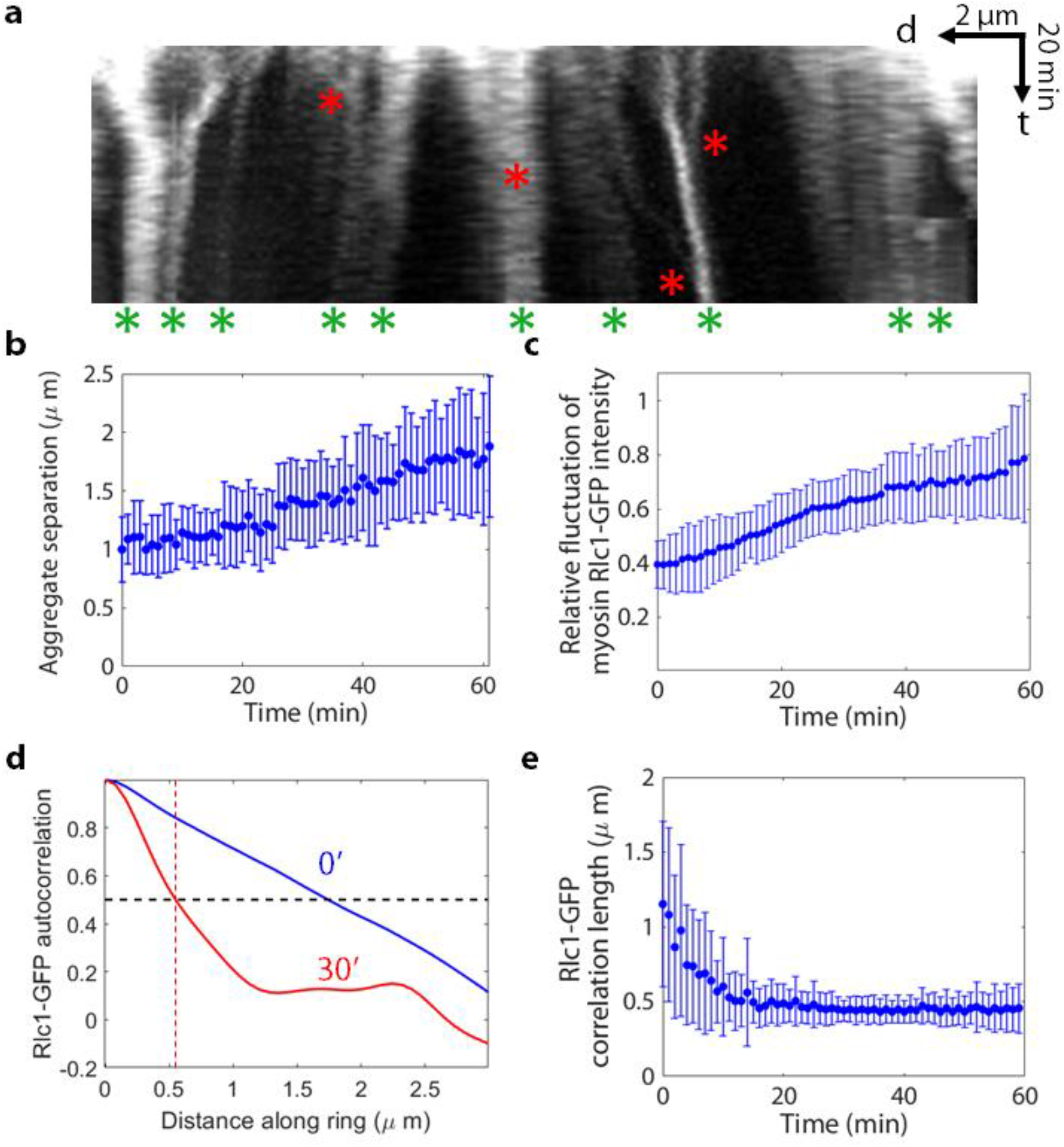
Myosin II aggregates hierarchically. (a) Top: kymograph of the intact ring of Fig. 1c. Time bar (t), 20 min; scale bar (d), 2 μm. Red asterisks: hierarchical aggregation events, when two aggregates merge to form an aggregate with more myosin II. Green asterisks: aggregates at the end of the observation time (60 min). (b) Mean separation between myosin II aggregates versus time, averaged over all aggregates in *n* = 10 rings. (c) Relative fluctuation of myosin Rlc1-GFP fluorescence intensity (ratio of s.d. to mean) versus time in the rings of (b). (d) Normalized Rlc1-GFP spatial correlations at the indicated times. The autocorrelation is the average of the product of deviations of the intensity from the mean at locations separated by a given distance around the ring, normalized to the value at zero separation. Red dashed line: the correlation length is the separation at which the correlation has decreased 50%. (e) Correlation length of Rlc1-GFP fluorescence intensity versus time, averaged over the rings of (b). (b,c,e) Error bars indicate s.d.

Myosin aggregated hierarchically. Neighboring aggregates moved toward one another and merged, and in some cases we observed merged aggregates themselves merging into higher order aggregates containing yet more myosin (Fig. 2a). Occasionally aggregates split into two parts. The progressive, hierarchical aggregation steadily increased the mean distance between neighboring aggregates from 1.00 ± 0.28 μm to 1.77 ± 0.56 μm over 60 min (Fig. 2b, *n* = 10, values are mean ± s.d.).

To quantify the punctateness we measured the fluctuation in myosin fluorescence intensity relative to the mean, which increased ~2-fold over 60 minutes (averaged over *n* = 10 rings, Fig. 2c). A measure of aggregate size is the half width of the spatial correlation function of the intensity around the ring (Fig. 2d), which decreased from 1.15 ± 0.55 μm to 0.46 ± 0.16 μm over 60 min, with most of the decrease in the first 20 min (Fig. 2e, values are mean ± s.d.).

### Molecularly explicit mathematical model of the contractile ring in cell ghosts

To explore the mechanisms underlying the observed myosin aggregation (Figs. 1, 2), we developed a 3D molecularly explicit mathematical model of the cytokinetic ring in *S. japonicus* cell ghosts, closely related to our previous model of the ring of the fission yeast *Schizosaccharomyces pombe*^11, 28^ (Figs. 3a, S1). For details, see Methods. Membrane-anchored formin Cdc12 dimers anchor actin filament barbed ends to the membrane. Actin filaments dynamically crosslinked by α-actinin are bound and pulled by myosin II according to a force-velocity relation set by the measured myosin II gliding velocity^29^. Myosin II is anchored to the plasma membrane in coarse-grained clusters of 8 Myo2 dimers^30^, with anchor drag coefficient in the membrane chosen to be consistent with the observed aggregation times.

**Figure 3.**
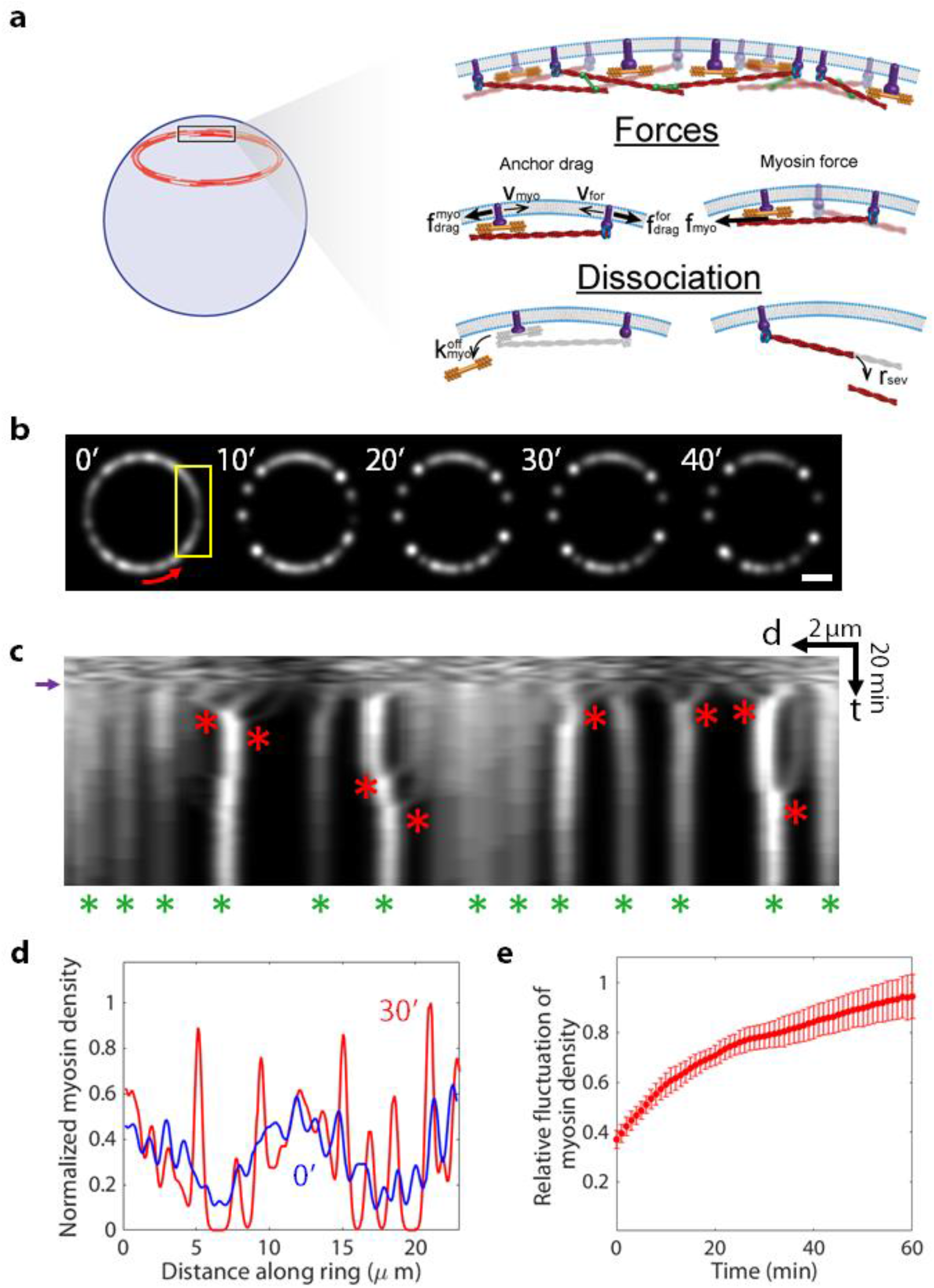
Molecularly explicit mathematical model of contractile rings lacking component turnover reproduces experimental hierarchical aggregation of myosin II. (a) Schematic of the cell ghost contractile ring model. Components do not turn over: association is absent, and dissociation is much slower than normal (see main text). Barbed-end anchored actin filaments bind and are pulled by membrane-anchored myosin II. Component motions are resisted by drag from the plasma membrane on membrane anchors. (b) Myosin-II distributions from simulations of the model. Simulated confocal fluorescence images are shown, using a point spread function of full width half maximum 600 nm similar to that of the microscope used for experimental images. The ring was run to steady state for 11 min under normal turnover conditions. Turnover was then abolished at *t* = 0. Red arrow: coordinate along ring. Scale bar: 2 μm. (c) Kymograph of the ring of (b). Time bar (t), 20 min; scale bar (d), 2 μm. After turnover is switched off (purple arrow), myosin II aggregates (red asterisks). Hierarchical aggregation leaves a diminished number of aggregates at the end of the simulation period (green asterisks). (d) Relative myosin II density versus distance around the ring of (b) at the indicated times. Densities are normalized so that the maximum density at 30 min is unity. (e) Simulated relative myosin II density fluctuations (ratio of s.d. to mean) versus time, averaged over *n = 10* rings. Error bars indicate s.d.

To model the turnover-free situation in *S. japonicus* cell ghosts, component association is entirely absent and the ring of radius 3.7 μm does not constrict (Fig. 3a). The dissociation rate of α-actinin was taken as the value measured for *S. pombe*^11, 31^, while that of myosin II was chosen to reproduce the slow decrease in total myosin-II fluorescence over time measured here (Methods and Fig. S2). Since rings in *S.japonicus* ghosts with phallacidin-stabilized actin filaments showed negligible actin loss^27^, we assumed formins do not dissociate. Rates of cofilin-mediated severing of actin filaments were set to reproduce the previously measured actin loss over ~40 min ^27^. The initial ring at the instant of ATP addition was assumed normal (Fig. 1c, d) and was generated by pre-equilibration ring for 11 min with normal component turnover, using component densities, actin filament lengths and association rates determined by previous measurements in *S. pombe*^30, 32^. In the following, images of simulated rings were convolved with a gaussian of width 600 nm to mimic the experimental microscope point spread function (see Methods).

### In simulated contractile rings myosin II aggregates hierarchically in the absence of turnover

In simulations of the model, the myosin distribution evolved in very similar fashion to that seen experimentally (Figs. 1,2). Within 5-10 minutes, isolated myosin II puncta had developed (Fig. 3b), presumably due to the lack of component turnover. Kymographs revealed that small initial aggregates emerged from the relatively homogenous initial distribution, some of which then merged into aggregates containing more myosin (Fig. 3c). Some of these aggregates underwent further rounds of merging, yielding aggregates with yet more myosin. As hierarchical aggregation proceeded, peaks in the myosin density profile sharpened (Fig. 3d) similar to the experimental fluorescence intensity (Fig. 1d). As a result, relative myosin density fluctuations increased ~ 2.5-fold over 60 mins (Fig. 3e), compared to the ~2-fold experimental increase in relative myosin fluorescence intensity (Fig. 2c).

The distinct myosin II aggregates that emerged after 5-10 min had a characteristic organization, with myosin II at the center and polarity-sorted outward-pointing actin filaments (filament barbed ends colocalizing with the myosin and distal pointed ends) (Fig. 4a). In a typical aggregation event, two such aggregates were pulled toward one another when the myosin in each aggregate engaged with the actin filaments hosted by the other aggregate and stepped toward the barbed ends. This occurred provided the aggregates were within a range of order the mean actin filament length of each other (Fig. 4a), taken as ~ 1.4 μm in our simulations from measurements on *S. pombe* rings^33^. Aggregates separated by more than this distance no longer affected each other, since filaments mediated the mechanical interaction. The end result of an aggregation was a similarly organized larger aggregate, with more myosin II and more polarity-sorted actin filaments.

**Figure 4.**
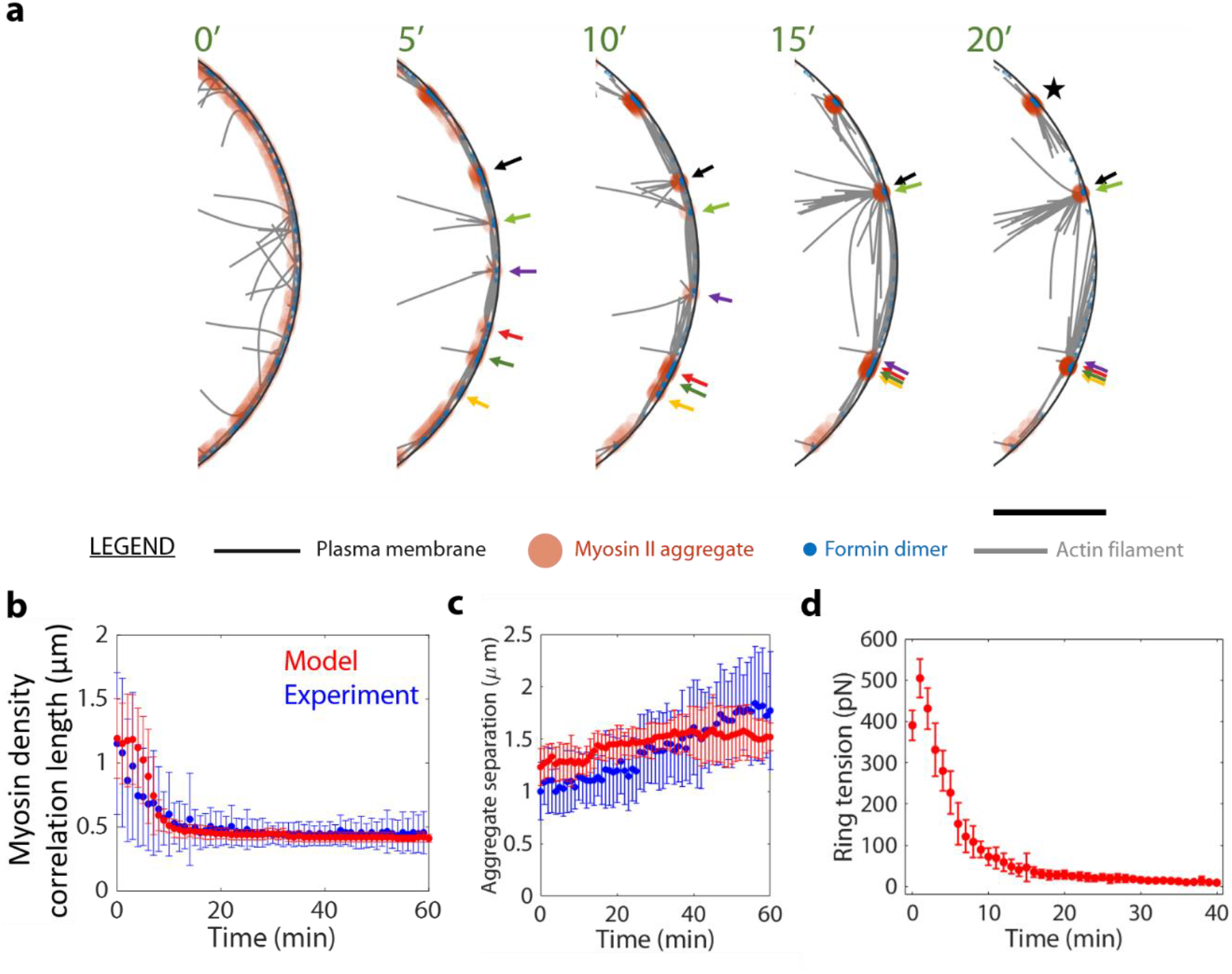
Actin filaments mediate mechanical communication among aggregates and set the aggregation range. (a) Section of the simulated cell ghost contractile ring of Fig. 3b (yellow box) at the indicated times. Myosin II aggregates progressively merge into more substantial aggregates (arrows; each color tracks a specific initial aggregate). The process continues until the large aggregates are no longer connected by actin filaments (e.g. aggregate labelled ★), when the ring fractures with tension loss. Formin dimer size and actin thickness not to scale. Model parameters as in Table S1. Scale bar: 1 μm. (b) Myosin II density correlation length versus time averaged over the simulated rings of Fig. 3e. Experimental data of Fig. 2e shown for comparison. (c) Mean myosin II aggregate separation versus time averaged over the simulated and experimental rings of Fig. 3e and Fig. 2b, respectively. (d) Mean ring tension versus time averaged over the simulated rings of Fig. 3e. ATP is added at time zero. Over 40 min the tension decreases to ~ 9 pN. Switching off actin filament elongation causes a very early transient tension increase. (b-d) Error bars are s.d.

These kinetics led to increasingly compact aggregates (Fig. 4b) separated by increasingly large distances (Fig. 4c). The size of a compact aggregate depended on the amount of myosin II it contained and the excluded volume interactions, reflecting steric interactions among myosin II molecules, which imposed a physically meaningful maximum density. The evolution of the correlation length and mean aggregate separation were both quantitatively consistent with experiment (Figs. 4b, c).

### Runaway myosin aggregation leads to ring fracture and tension loss

With time, the unopposed myosin aggregation destroyed the structural integrity of simulated rings and virtually abolished the ring tension. After ~ 20 min rings lost mechanical connectivity (Movie 1) and ring tension decreased from ~ 500 to 9 pN over 40 min (Fig. 4d), compared to the ~ 400 pN measured in contractile rings of *S. pombe* protoplasts^11^.

### Runaway aggregation is rescued by myosin II turnover but not by actin turnover

Our results suggest that in normal cytokinetic rings turnover of myosin and actin protect the organization against contractile instability (Fig. 5a). We next asked if prevention of runaway aggregation required turnover of one or both of these components. We ran cell ghost simulations with either myosin-II turnover or actin turnover restored to normal. Myosin II turnover alone (with the turnover rate measured in *S. pombe*, .026 s^-1 31^) prevented aggregation and rescued the ring organization (Figs. 5b,d). However, restoring actin turnover (with turnover rates consistent with experimental measurements in *S. pombe*, see Methods) failed to rescue rings from catastrophic aggregation (Fig. 5c). In the latter case, compared to cell ghosts late stage aggregates had a larger separation (Fig. 5e), since actin turnover maintained longer filaments that allowed myosin aggregates to communicate over greater distances, and the rate of tension loss was lower (Fig. S2c).

**Figure 5.**
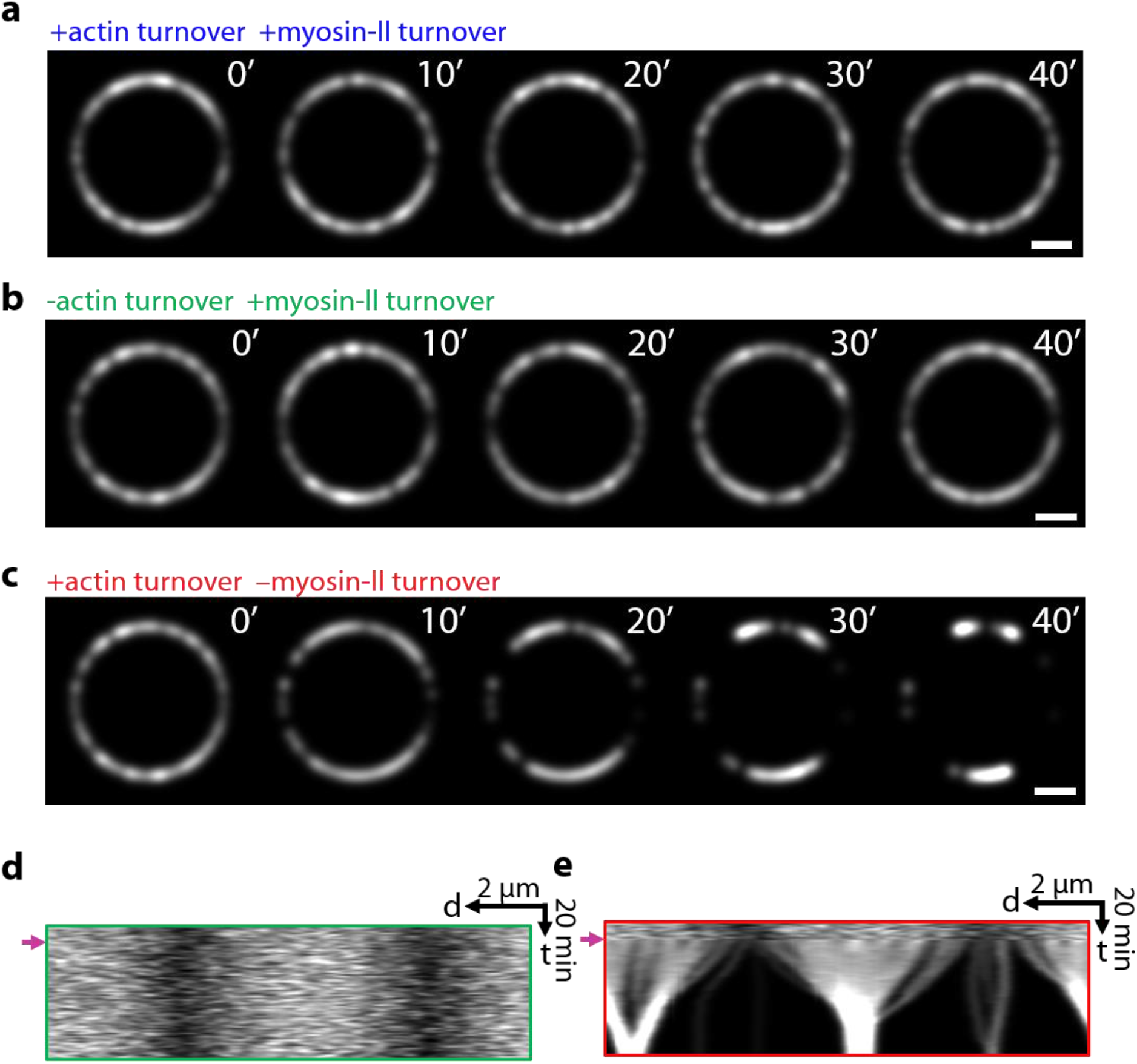
Runaway aggregation is rescued by myosin II turnover but not by actin turnover. (a-c) Simulated confocal fluorescence micrographs of myosin II in cell ghosts with either myosin-II turnover restored (b), or actin turnover restored (c), or turnover of both components restored (a). Scale bar: 2 μm. (d,e) Kymographs of the rings of (b,c), respectively. Arrows indicate the instants when the respective turnover conditions were implemented.

### Myosin diffusion is too weak to prevent runaway aggregation due to contractile instability

The present work implicates myosin II turnover as the mechanism that controls contractile instability in cells. It has been suggested that myosin II diffusivity serves this purpose, by smoothing small scale density variations that characterize the instability^14, 25^. To test this proposal we ran simulations of contractile rings without turnover as previously, but now the membrane-anchored myosin II clusters could diffuse laterally in the membrane with diffusivity *D*.

We tested diffusivities in the range 10^-5^ ≤ *D* ≤ 10^-2^ μm^2^s^-1^ (Fig. 6). Myosin aggregation was not prevented by diffusion, except for the highest values tested *D* ~10^-2^μm^2^ s^-1^. This value is almost three orders of magnitude greater than measured diffusivities of membrane-bound myosin II in fission yeast contractile rings, *D*~2 × 10^-5^μm^2^s^-1 34^. We conclude that physiologically relevant myosin diffusion is far too weak to oppose contractility-induced aggregation.

**Figure 6.**
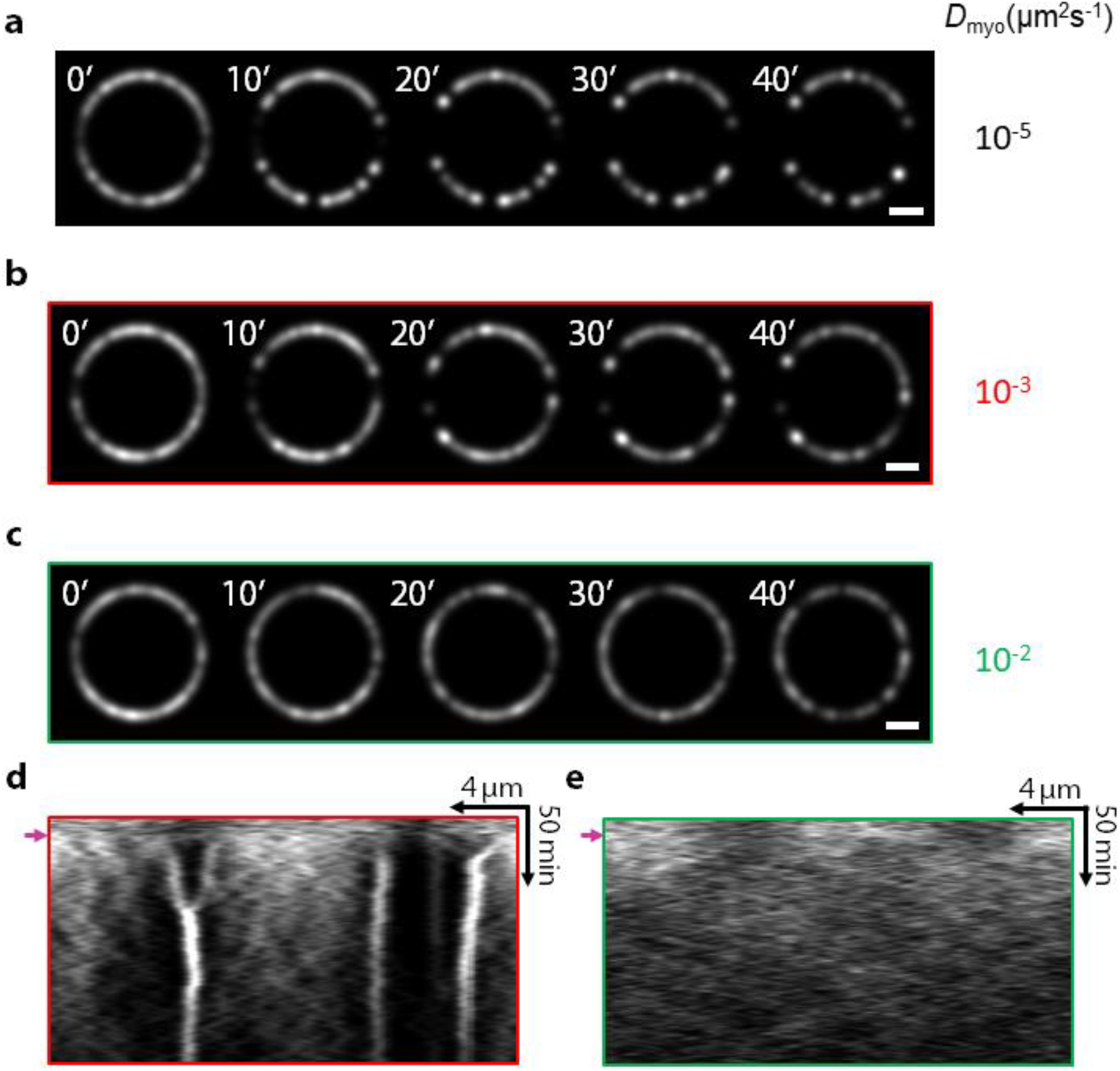
Myosin II diffusion does not prevent runaway aggregation due to contractile instability. (a)-(c). Simulated confocal fluorescence micrographs of myosin II in cell ghost contractile rings that lack turnover, with lateral diffusion of myosin II aggregates in the plasma membrane implemented. Three diffusivities were simulated, *D*_myo_ = 10^-5^, 10^-3^ and 10^-2^ μm^2^s^-1^. Myosin diffusion rescues the normal myosin II distribution only for the largest diffusivity, a value ~500 fold larger than physiological values measured in fission yeast *S. pombe^34^*. Scale bar: 2 μm. (d,e) Kymographs of the rings of (b) and (c) respectively. Arrows indicate instants at which turnover is switched off.

## Discussion

The cytokinetic contractile ring and other actomyosin machineries require mechanisms to intercept contractile instability (Fig. 1a), but these have been difficult to identify since the full course of the instability is rarely observed. Here, we observed the full instability in isolated contractile rings in cell ghosts. In a hierarchical process, myosin II aggregated into puncta that merged with one another, some of these larger aggregates merging into yet larger aggregates, and so on (Fig. 2a). Aggregation progressively increased the myosin per aggregate and the aggregate separation, up to ~ 1-2 μm (Fig. 2b). We observed no greater separations, suggesting that this separation effectively terminated the process, consistent with simulations which identified the maximum separation as the mean actin filament length (Fig. 5).

Here, hierarchical aggregation is driven by active myosin-mediated contractile forces. It is of interest to compare this with analogous systems where the driving forces are passive particle-particle adhesive interactions, well known in contexts such as aggregation of atmospheric pollutant particles or colloidal suspensions^35^. In these cases larger aggregates are effectively more adhesive due to their greater surface area, which can lead to hierarchical kinetics dominated by ever-larger aggregates^36^.

We reasoned that the uncontrolled aggregation was caused by the absence of component turnover. Since cell ghosts lack cytoplasm, association of myosin II, actin and other components is absent and component dissociation rates are severely reduced. Turnover would tend to restore homogeneity to the ring, so its absence could explain the runaway aggregation. Molecularly explicit simulations corroborated this interpretation (Figs. 3–6). Simulations of cell ghost rings lacking turnover quantitatively reproduced the experimental hierarchical myosin aggregation. Aggregates emerged from local stochastic myosin density peaks that drew in myosin and actin filaments (Fig. 1a). Late stage aggregates had a myosin II core and outward pointing polarity-sorted actin filaments. Interestingly, some late aggregates had an aster structure, with a relatively compact myosin core and actin filaments pointing radially outward in multiple directions (Fig. 4a), similar to *in vitro* actomyosin systems which evolved to a final state consisting of morphologically quiescent asters with polarity sorted actin filaments^37^. Other late aggregates had a more extended myosin II core, with longitudinally oriented actin filaments emerging at the edges (Fig. 4a).

Hierarchical aggregation, in which aggregates repeatedly merged into yet larger aggregates, continued until the aggregate separation exceeded the typical actin filament connector length when aggregates could no longer communicate mechanically. The contractile instability had then run its full course, reaching a final state consisting of unchanging aggregates, similar to the final state observed *in vitro*^37^. At this stage, ring fracture and tension loss occurred (Figs. 3,4).

By contrast, in simulations of rings in normal cells with turnover, myosin reached a steady state punctateness with moderately sized aggregates (Fig 5a). Myosin II turnover was the key process averting runaway aggregation, by replacing large myosin aggregates with more homogenously distributed moderately sized aggregates, before hierarchical aggregation could run its catastrophic course. Aggregation was prevented even with myosin II turnover alone, but no actin turnover, while actin turnover alone was insufficient (Figs. 5b,c).

Our previous experimental work emphasized the role of actin turnover for normal organization and function of contractile rings^27^. Myosin II aggregation in *S.japonicus* cell ghosts was abolished by treatment with the myosin II inhibitor blebbistatin, and stabilization of actin with Jasplakinolide suppressed formation of small myosin puncta and increased the fraction of rings that shortened. These rings appeared to fully constrict, but broad myosin II aggregates were apparent in these shortening rings, similar to our simulations with restored actin turnover (compare Figs. 3b and 5c). The shortening segments in these rings likely have zero tension, since a mathematical model^28^ of the analogous phenomenon seen experimentally^26^ in *S. pombe* fission yeast ghosts accurately reproduced the observed shortening rates and showed that rings shortened due to zero tension detached segments being reeled in by myosin II against zero load. For proper cytokinesis and contractile ring homeostasis in normal cells, turnover of both myosin II and actin likely both contribute in complimentary ways.

We conclude that rapid myosin turnover in contractile rings prevents runaway aggregation and tension loss due to large scale contractile instabilities. Contractile instability also affects the smallest scales since, if unopposed, it would aggregate myosin II into clumps of arbitrarily high density (see Fig. 1a). Simulated aggregate sizes without turnover were set by the amount of myosin II in the aggregates, excluded volume interactions and local polarity sorting (Fig. 4a). Our results suggest that in the cytokinetic ring and other actomyosin machineries, small scale contractile instability is likely controlled by several small scale effects. These include excluded volume interactions due to steric intermolecular interactions that impose packing limits, small scale actomyosin polarity sorting that suppresses contractility^37^, and other local interactions such as myosin II minifilament stacking in animal cells^38^.

It is often assumed that myosin diffusion provides this small scale control. Active gel models of actomyosin cortices comonly assume a diffusive contribution to the myosin motion, which helps suppress actomyosin instability that would otherwise cause uncontrolled short scale density blow up^14, 25^. To be effective, diffusivities of order *D*~0.01 μm^2^s^-1^ or larger are invoked. However, measured myosin II diffusivities in contractile rings in fission yeast^34^ are ~ 2 × 10^-5^ μm^2^s^-1^, ~3 orders of magnitude smaller, and were reported unmeasurably small in *C. elegans* rings^15^. Thus, myosin diffusivity is presumably not responsible for controlling actomyosin contractile instability, and its inclusion in models is a numerical convenience that misrepresents at least the small scale behavior. We confirmed this in simulations with added myosin cluster diffusivity. A diffusivity ~500 times larger than the experimental value was required to prevent uncontrolled aggregation (Fig. 6).

Overall, our results suggest that in actomyosin assemblies a balance of turnover and small scale effects regulate contractile instability on large and small scales, respectively (Fig. 7). We suggest contractile instability serves the useful function of assembling myosin II into force-generating aggregates, regulated by turnover which imposes a time limit on the contractile instability-driven aggregation. The steady state myosin aggregate size is set by the turnover time, together with local effects that dictate a maximum density of actomyosin components. The turnover-imposed time limit is sufficient for powerful myosin aggregates to be built, but short enough to prevent runaway aggregation and disastrous organizational disruption, tension loss and ring fracture (Fig. 7).

**Figure 7.**
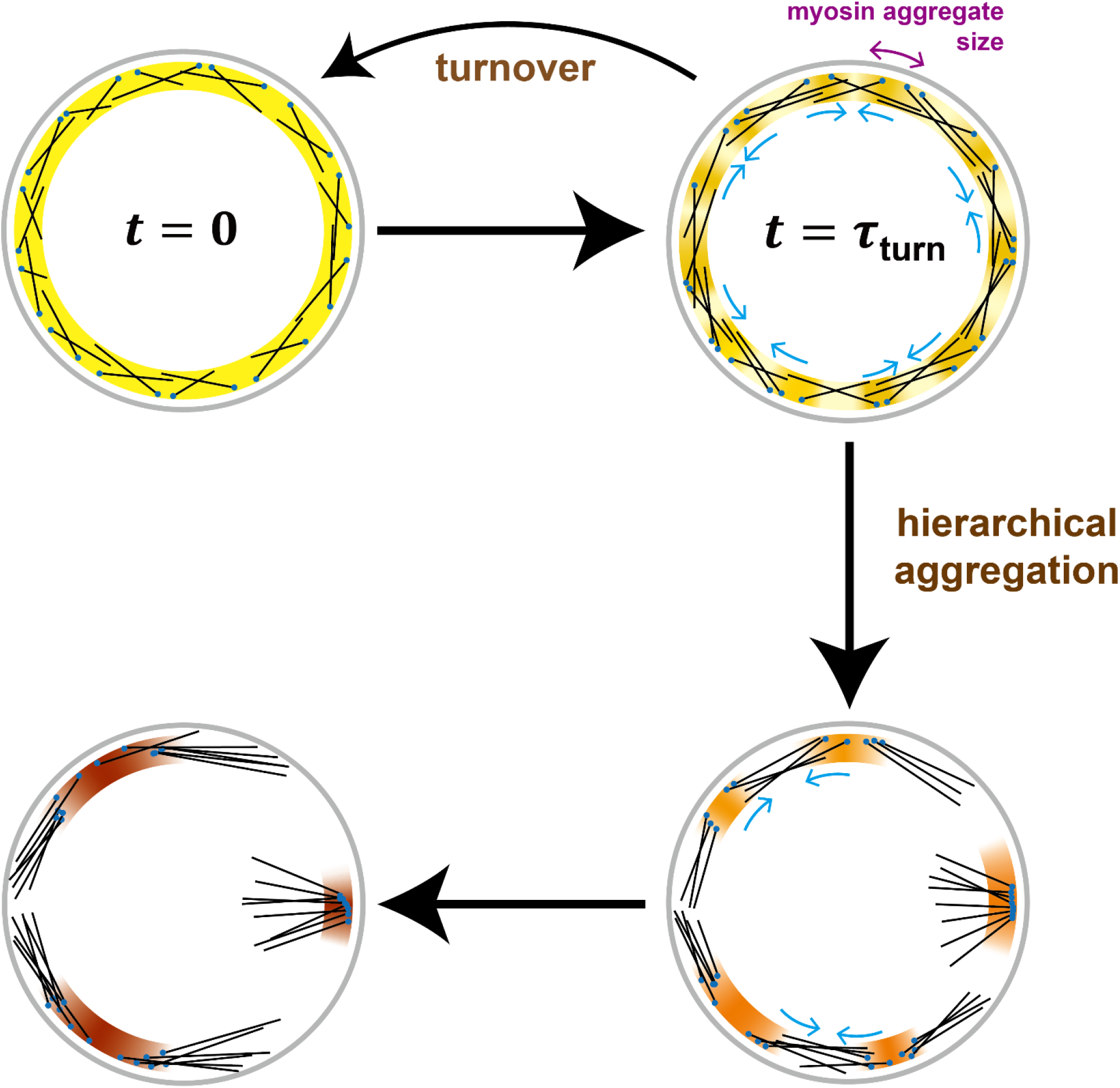
Turnover regulates contractile instability to set myosin II aggregate size and prevent runaway aggregation. Model of turnover-regulated contractile instability. Contractile instability progressively increases punctateness in the contractile ring and other actomyosin machineries. Given a homogenous initial distribution (*t* = 0), in normal cells component turnover tends to restore homogeneity by intercepting the instability on the turnover timescale *τ*_turn_, setting a functional steady state myosin II aggregate size. Without turnover contractile instability progresses unopposed, separating the aggregates and hierarchically merging them into yet bigger aggregates with increased separation. Late stage aggregates have a myosin II core and outward pointing polarity-sorted actin filaments, with a size governed by local excluded volume interactions that cap the density, and polarity-sorting that effectively switches off contractility. Once the aggregate separation exceeds the actin filament length aggregates can no longer communicate mechanically, and contractile instability has run its full course (bottom left).

## CODE AVAILABILITY

Simulations and quantitative analysis of images were performed in MATLAB. The source code is available upon request.

## ACKNOWLEDGMENTS

We thank Yutao Li for assistance with image processing, Dong An for assistance with figures, and Zach McDargh for helpful discussions. This work was supported by National Institute of General Medical Sciences of the National Institutes of Health under award number R01GM086731 to B.O’S. The content is solely the responsibility of the authors and does not necessarily represent the official views of the National Institutes of Health.

## AUTHOR CONTRIBUTIONS

B.O’S. designed the research; B.O’S., S.T., and S.W. performed the mathematical modeling; T.G.C. and J.H. performed the experiments; S.T. and S.W. analyzed the experimental data; B.O’S. and S.T. wrote the paper with contributions from all authors.

## Methods

### 1. S. japonicus protoplast and cell ghost preparation and ATP-dependent contraction of rings in cell ghosts

*S. japonicus* cell ghosts were prepared using protocols published previously^39, 40^. A brief summary is given below. For strains, see ref. 27. *S. japonicus* cells were cultured until midlog phase using a rich YEA medium, and cell walls were digested using lytic enzymes Lallzyme MMX (Lallemand; ref. 41) for protoplast preparation. Protoplasts were then washed and recovered in sorbitol-containing medium. Following ring assembly, protoplasts were washed once with wash buffer (20 mM Pipes-NaOH, pH 7.0, 0.8 M sorbitol, 2 mM EGTA, and 5 mM MgCl_2_) and permeabilized with isolation buffer (50 mM Pipes-NaOH, pH 7.0, 0.16 M sucrose, 50 mM EGTA, 5 mM MgCl_2_, and 50 mM potassium acetate) containing 0.5% NP-40 to obtain cell ghosts. Ghosts were washed twice with reactivation buffer (0.16 M sucrose, 5 mM MgCl_2_, 50 mM potassium acetate, and 20 mM MOPS-NaOH, pH 7.0; pH adjusted to 7.5) after permeabilization. Both the isolation and reactivation buffers were cooled to 4°C. The isolation and washing steps were performed on ice. Equal volumes of cell ghosts and reactivation buffer containing 1 mM ATP (A6559; Sigma-Aldrich) were mixed to induce ATP-dependent actomyosin ring contraction. It takes at least 1-2 mins after the addition of ATP to mount the ghosts and to find those appropriate for imaging. Thus, time 0 in figures represent the instant at which imaging commences.

### 2. Sample preparation for microscopy imaging

An equal volume of reactivation buffer containing 1 mM ATP was added to cell ghosts and imaging was performed in an Ibidi μ-Slide eight-well glass-bottom dish(#80827). To prevent water evaporation during imaging, an adhesive film membrane was used to seal all imaging dishes.

### 3. Spinning-disk confocal microscopy

Image acquisition was performed using Andor Revolution XD spinning disk confocal microscopy. The microscope was equipped with a Nikon Eclipse Ti inverted microscope, Nikon Plan Apo Lambda 100×/1.45-NA oil-immersion objective lens, a spinning-disk system (CSU-X1; Yokogawa Electric Corporation), and an Andor iXon Ultra EMCCD camera. Andor IQ3 software was used to acquire images at 80 nm/pixel. Laser lines at wavelengths of 488 or 561 nm were used to excite the fluorophores. Most images were acquired with *z*-step sizes of 0.2, 0.3, or 0.5 μm, and at various interval times.

### 4. 3D molecularly explicit model of the cytokinetic ring in *S. japonicus* protoplasts

We developed a fully 3D and molecularly explicit mathematical model of the *S. japonicus* cytokinetic ring. Formin Cdc12 dimers anchored to the membrane nucleate and grow actin filaments that bind myosin-II, assumed membrane-anchored from previous studies. In this section, we describe the ring before turnover is abolished.

#### 4.1. Ring geometry

The ring lies on a cylinder, at the inner surface of the plasma membrane. Ring components bind with uniform probability in a binding zone of width 0.2 μm^32, 42^ on the plasma membrane, thus determining the width of the ring. The thickness of the ring in the direction perpendicular to the plasma membrane is decided by simulation dynamics. We used a ring radius of 3.7 μm as the rings we observe here have a mean length up to ~22-25 μm.

#### 4.2. Components in the ring

We describe actin filaments as inextensible but flexible polymers with bending modulus *κ* = *k_B_Tl_p_* where the persistence length *l_p_* = 10 μm^43, 44^. In the simulation, we keep track of every 37^th^ actin subunit on a filament, with spacing 0.1 μm from each other, as in our earlier works^28, 42^. The subunits at the barbed end of each filament represent formins, which are anchored to the plasma membrane and subject to anchoring drag.

Myosin-II in the ring is described as clusters containing 16 myosin heavy chains^28^, and are anchored to the plasma membrane and subject to anchoring drag^42^. A cluster binds and pulls on actin filaments within its capture radius *r*_myo_ = 80 nm measured from the center of the cluster^45^.

Alpha-actinin crosslinkers are modeled as springs with rest length 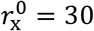 nm and spring constant *k*_x_ = 25 pN/μm, connecting pairs of actin subunits (only those that we keep track of) on different filaments that are within distance 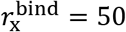 nm from each other.

#### 4.3. Forces in the ring

##### Binding of actin filaments to myosin clusters

A myosin-II cluster α binds and exerts a capture force 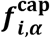 on every actin subunit *i* within its capture radius *r*_myo_, implemented as a spring with spring constant *k*_cap_ = 400 pN/μm and zero rest length that connects the center of the cluster ***r_α_*** to the actin subunit located at ***r_i_***. In order not to interfere with the pulling force of the myosin, we decompose this force along and perpendicular to the actin filament, and only the latter is actually applied.

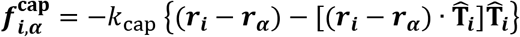

where 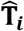 is the unit tangent vector of the actin filament at subunit *i*.

##### Force-velocity relation of myosin

Myosin-II cluster α pulls on bound actin subunits *i* with a force tangent to the filament

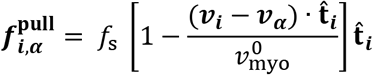

where 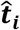 is the unit vector from the (*i*-1)^th^ subunit to the *i*^th^ subunit. This force decreases linearly with the velocity that the myosin-II cluster moves toward the barbed end (***v_i_ – v_α_***), and is equal to the stall force *f*_s_ = 4 pN or zero when the velocity is equal to zero or the load-free velocity 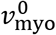 respectively. When one myosin-II cluster pulls on *n*_fil_ > 10 actin filaments, the stall force is lowered to a value *f*_s_ = (4 pN) × (10/*n*_fil_).

##### Excluded volume interactions

Myosin-II clusters repel other clusters located within a distance *d*_myo_ = 45 nm, with an elastic force that increases linearly as the distance decreases. The elastic constant 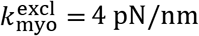. The value of the distance was chosen so that the aggregates from the simulation and experiment had a similar morphology, which is a mix of broad and point-like aggregates.

##### Tension in actin filaments and ring tension

Tension is implemented as pairwise attractive force of magnitude 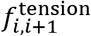 between adjacent actin subunits labelled by *i* and *i* + 1. The constraint force ensures that the subunit separation is maintained at 0.1 μm (see subsection 1.2 above). The force is implicitly set by the dynamics, and in practice is calculated together with component velocities. See “Computation scheme” of subsection 1.5 below. The sum of filament tension forces exerted across each ring cross section were obtained and averaged over all cross-sections to obtain ring tension at every time instant.

##### Bending forces of actin filaments

Following the scheme in Ref. ^46^, first the bending energy of one filament is calculated as

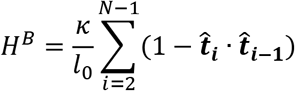

The filament has *N* subunits, each separated by *l*_0_ = 0.1 μm, and a bending modulus *κ*. The bending force is given by –(*∂H*^B^)/(*∂**r_i_***), the negative derivative of *H^B^* with respect to the coordinates of the subunits.

##### Crosslinking

Crosslinkers exert a spring force 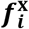 between actin subunits. See section “Ring components” for details.

##### Ring component confinement

This constraint is implemented as an elastic restoring force with elastic constant *k*_mb_ = 20 pN/μm pointing towards the origin that is activated when any ring component is greater than R away from the origin, where R is the radius of the protoplast.

##### Normal anchoring forces

Formin Cdc12 dimers and myosin-II clusters are anchored to the plasma membrane^30^, and an anchoring force normal to the plasma membrane maintains their radial coordinate at *R* – *d*_for_ and *R* – *d*_myo_ respectively, where *R*, *d*_for_, *d*_myo_ are the radius of the protoplast, and distances of formin and myosin from the membrane respectively. The magnitude of this force is implicitly set by the dynamics, and in practice is calculated together with component velocities. See section “Computation scheme” below. The values of *d*_for_ and *d*_myo_ are 30 nm and 80 nm respectively, based on super-resolution measurements^30^.

##### Tangential anchoring forces due to membrane drag

As all formin Cdc12 dimers and myosin-II clusters are membrane anchored, they experience drag forces 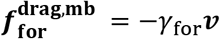 and 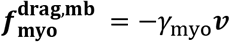 respectively.

##### Cytosolic drag forces

Drag forces for actin subunits in the cytosol are set by their velocities and the drag coefficient *γ*_act,proto_ = 0.2 pN · s · μm^-1 42^. Cytosolic drag forces for myosin-II clusters and formin Cdc12p dimers are neglected as they are much smaller than the membrane drag force.

##### Stochastic forces

A myosin-II cluster *α* of the simulations in Fig. 6 has a diffusivity *D* due to a newly added stochastic, zero-mean force 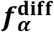. The correlation of the force is given by 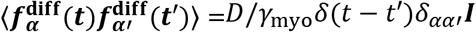 where *γ*_myo_ and ***I*** are the drag coefficient along the membrane and the identity matrix respectively.

#### 4.4. Turnover of ring components

Formin Cdc12p dimers, myosin-II clusters and α-actinin crosslinks unbind the ring at rate 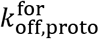, 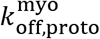 and 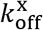, respectively. In addition, α-actinin unbinds the ring if the two actin subunits that it crosslinks are separated by > 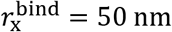. Formin Cdc12 dimers and myosin-II clusters bind the plasma membrane in a binding zone of width 0.2 μm. Alpha-actinin crosslinks bind the ring with equal probability between any pair of actin subunits within 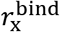 unless this pair has already been crosslinked. Component binding rates are chosen to maintain steady state of component densities (Table S1). Crosslinker binding rates are tuned to achieve a steady-state cross-linker density of 25 μm^-1 32^.

Actin is polymerized by formin at *v*_pol_ = 70 nm/s, and subject to severing by ADF-cofilin at rate *r*_sev,proto_ = 1.8 μm^-1^min^-1^, at a random location along the filament with uniform probability^42^. Once severed, the portion from the severing point to the pointed end is removed from simulation.

Mean actin filament length in simulations is ~ 1.4 μm, and is set by formin density, filament growth rate, and filament severing rate. (1) Previously measured formin densities are ~ 15 per μm (Table S1). (2) Electron micrographs showed ~ 20 filaments in the cross section of the *S. pombe* ring^47^. (3) Upon addition of the actin monomer sequestering drug Latrunculin A to intact cells, in 60 s, ~90% of cells lost their rings^48^. The growth and severing rates *v*_pol_ and *r*_sev_ were obtained as best-fit parameters to reproduce (2) and (3) above.

The initial experimentally measured Rlc1-GFP fluorescence distribution had a correlation length ~1.0 μm (Fig. 2e). In our simulations we used a simple measure to reproduce these statistics. We divided the perimeter of the simulated ring into four equal sectors and chose the association kinetics such that incoming myosin clusters had a preference to bind to two non-neighboring sectors. This generated steady-state rings with a correlation length of myosin density at *t = 0* similar to that seen experimentally before turnover was abolished (Fig. 4b). Irrespective of whether or not we used such a spatial turnover bias to set up the initial state of the ring, we observed the same hierarchically aggregating myosin behavior once turnover was switched off.

#### 4.5. Simulation of the model

##### Initial configuration of the ring

In order to mimic the experimental situation, where the ring in the protoplast is allowed to fully assemble before the protoplast is permeabilized, we set our time zero to the steady state of the ring.

The ring is initially a 0.2 μm wide bundle composed of myosin-II clusters and actin filaments that have a clockwise or anti-clockwise orientation with equal probability. Their barbed ends (formin Cdc12p dimers) and myosin-II clusters are randomly distributed in a band-like zone 0.2 μm wide on the plasma membrane, with uniform probability per area. 11 minutes of ring dynamics is simulated in order for the ring to reach steady state.

##### Numerical scheme

Given the ring configuration at any time step, we numerically solve the linear system of force balance equations

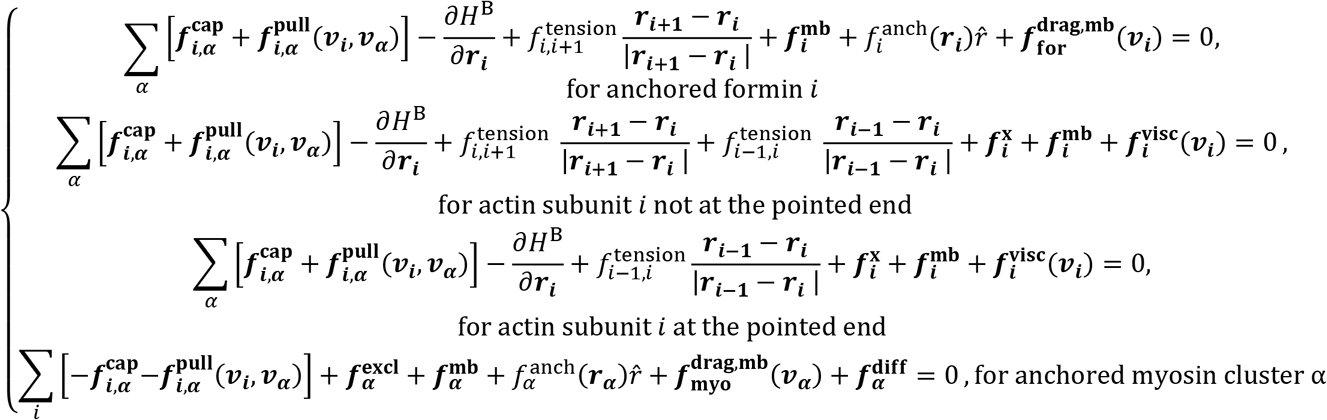

and the constraint equations

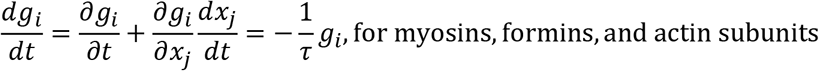

for the unknown variables {***v_i_***}, {***v_α_***}, and the Lagrange multipliers 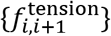, 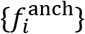 and 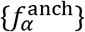 for each ring component. The constraints {*g_i_*({*x_j_*}) = 0} represent the anchoring of formins and myosins to the membrane, and the maintenance of distance between neighboring actin subunits (see subsection 1.3 above). The Euler method with a time step Δ*t* = 0.01 s and *τ* = 10Δ*t* is used to evolve the system. The scheme was adapted from Witkin et al.^49^. At each time step, components are added to and removed from the simulation according to turnover dynamics.

To optimize simulation running time and enable larger timesteps, we added a pairwise drag force ***f*** = –*γ*_a_**Δ*v*** between every pair of neighboring actin subunits and between an actin subunit and its bound myosin cluster, in order to suppress spurious oscillations. Here, **Δ*v*** is the relative velocity in each pair and the artificial drag coefficient *γ*_a_ = 4 pN · s/μm. No net force was added to the simulation as this is a pairwise interaction.

### 5. 3D molecularly explicit model of the cytokinetic ring in permeabilized *S. japonicus* protoplasts (ghosts)

We modified the protoplast model, in order to describe the ring in permeabilized *S. japonicus* protoplasts. Ring components no longer bind to the ring, and unbinding dynamics are slowed down. The fraction of actin in the ring in permeabilized protoplasts does not decrease with time upon treatment with ATP and phallacidin^27^. This implies that formin (bound to the barbed end of actin filaments) unbinds the ring with negligible rate, 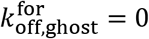. We tune cofilin-mediated severing rate *r*_sev,ghost_ in permeabilized protoplasts to reproduce the fraction of actin remaining in the ring at 40 min after treatment with ATP alone (Table S1). Actin polymerization is absent, *v*_pol_ = 0. We tuned the myosin-II cluster off rate 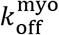 to be consistent with our measurements of the time course of Rlc1-GFP intensity (Fig. S2a). Here, we have interpreted the decrease in myosin-II intensity over time as due to myosin-II loss from the ring. However, we found that myosin II aggregation kinetics were qualitatively unchanged even when we assumed there was no myosin II loss whatsoever, by setting 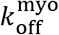 to zero (Fig. S2b). (In this interpretation, the decrease in Rlc1-GFP intensity of Fig. S2a would be attributed to bleaching.) Cross-linker binding rates are set to zero. For the rings of Fig. 5, along with cross-linker turnover, either actin or myosin turnover processes were abolished as indicated.

### 6. Image Analysis

We analyzed three-dimensional confocal micrographs of rings in cell ghosts in ImageJ^50^. After isolating a 3D volume containing the ring, we identified a best-fit plane where the maximum amount of Rlc1-GFP fluorescence was present. We then rotated the image such that the Z axis of the rotated 3D image is parallel to the normal to the best-fit plane, and performed a sum intensity projection to get a 2D image. We performed background subtraction using the rolling ball technique with a radius of 50 pixels. We measured the total fluorescence in a rectangular region containing the ring (Fig. S2a). We used contours of finite thickness coincident with the ring to generate kymographs using the “max” option in the KymographBuilder plugin. Kymographs were analyzed in MATLAB. Aggregates were identified as peaks in the intensity at each time instant in kymographs. We only indicated aggregation events in Figs. 2a and 3c which were flagged by the algorithm, with the exception of the third red asterisk from the left in Fig 3c which we judged to be a valid aggregation event despite not being flagged by the algorithm. Brightness and contrast of the entire image were adjusted for presentation purposes for Fig. 1c and Fig. 2a.

Images of simulations were generated by convolving myosin positions with a gaussian with a full-width half-maximum of 600 nm to mimic the experimental microscope point spread function (PSF). Rings in our experimental images are oriented arbitrarily in 3D space and thus are affected by the small PSF width in the X-Y plane, and the comparatively larger PSF widths in the X-Z and Y-Z planes.

---

**Movie 1.** Simulated cell ghost contractile ring of Fig. 3b at the indicated times. Turnover was abolished at t = 0. Formin dimer size and actin thickness not to scale. Scale bar: 1 μm.

