## Supplementary Materials for "Myosin turnover controls actomyosin contractile instability"

### Supplementary Figures

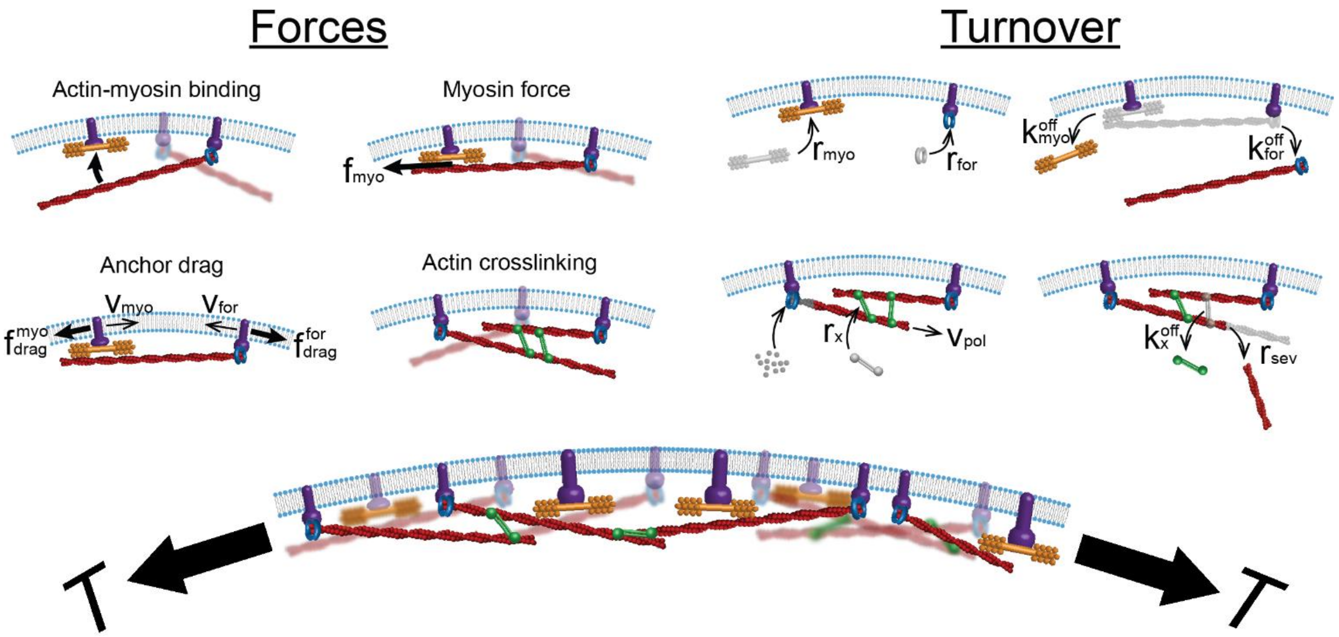


**Supplementary Figure 1. 3D model of the cytokinetic ring in fission yeast protoplasts (components not to scale).** See Supplementary Table 1 for component amounts and key parameters. Actin filaments are anchored to the plasma membrane at barbed ends by formin Cdc12, and bind membrane-anchored myosin-II that pulls them according to a linear force-velocity relation. Component motions are resisted by drag from the cytoplasm and the plasma membrane (on anchors only). These components constantly turnover: actin is polymerized by formin Cdc12 dimers, dissociates by unbinding with formins, and by stochastic cofilin-mediated filament severing; myosin-II clusters, formin Cdc12 dimers and α-actinin crosslinks constantly bind and unbind the ring. In cell ghosts, components no longer bind the ring, and unbinding dynamics are slowed down (see Table S1).

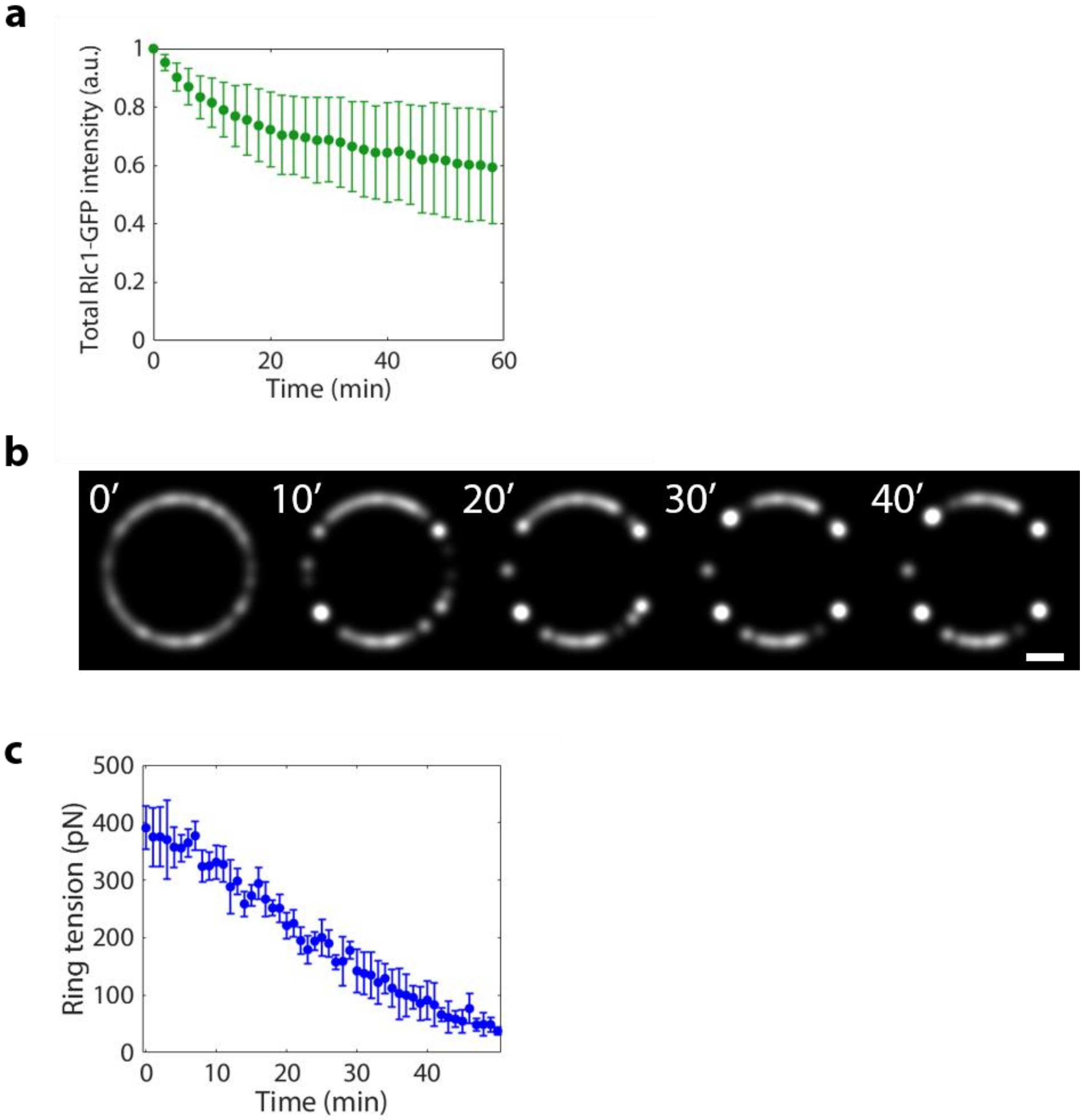


**Supplementary Figure 2. Myosin aggregation in simulations with no myosin dissociation or with actin turnover but no myosin turnover.** (a) Mean total Rlc1-GFP fluorescence intensity along the ring versus time, averaged over the rings of Fig. 2b. Error bars indicate s.d. (b) Simulated confocal fluorescence images of myosin-II distributions from simulations of the model where there was no myosin loss after turnover was abolished. Figure preparation and simulation protocol similar to that used in Fig. 3b. Scale bar: 2 μm. (c) Mean ring tension versus time averaged over simulated rings with only actin turnover restored (*n* = 5 rings, error bars are s.d.). ATP is added at time zero.

**Supplementary Table 1. Key parameters values for simulations of rings in *S. Japonicus* cell ghosts.**

| Parameter | Meaning | Value | | Legend |
|---|---|---|---|---|
| | | in protoplasts | in cell ghosts | |
| $\rho_{\mathrm{for}}$ | Initial mean formin dimer density | $15\ \mu\mathrm{m}^{-1}$ | | (A) |
| $\rho_{\mathrm{myo}}$ | Initial mean myosin cluster density | $18.75\ \mu\mathrm{m}^{-1}$ | | (B) |
| $r_{\mathrm{sev}}$ | Actin filament severing rate by cofilin | $1.8\ \mu\mathrm{m}^{-1}\mathrm{min}^{-1}$ | $0.0409\ \mu\mathrm{m}^{-1}\mathrm{min}^{-1}$ | (C) |
| $k_{\mathrm{off}}^{\mathrm{for}}$ | Off rate of formins | $0.023\ \mathrm{s}^{-1}$ | 0 | (D) |
| $k_{\mathrm{off}}^{\mathrm{myo}}$ | Off rate of myosin clusters | $0.026\ \mathrm{s}^{-1}$ | $2.2\ \times 10^{-4}\ \mathrm{s}^{-1}$ | (E) |
| $k_{\mathrm{off}}^{\mathrm{x}}$ | Off rate of α-actinin | $3.3\ \mathrm{s}^{-1}$ | | (F) |
| $f_{\mathrm{s}}$ | Myosin stall force per filament | $4\ \mathrm{pN}$ | | (G) |
| $v_{\mathrm{myo}}^{0}$ | Myosin load-free velocity | $0.24\ \mu\mathrm{m/s}$ | | (H) |
| $r_{\mathrm{myo}}$ | Capture radius of myosin-actin binding | $80\ \mathrm{nm}$ | | (I) |
| $l_{\mathrm{p}}$ | Persistence length of actin | $10\ \mu\mathrm{m}$ | | (J) |
| $\gamma_{\mathrm{myo}}$ | Membrane anchor drag coefficient of myosin cluster | $9.36\ \mathrm{nN}\cdot\mathrm{s}/\mu\mathrm{m}$ | | (K) |
| $\gamma_{\mathrm{for}}$ | Membrane anchor drag coefficient of formin dimer | $1.9\ \mathrm{nN}\cdot\mathrm{s}/\mu\mathrm{m}$ | | (L) |

(A) Ref. 1.

(B) Using 16 heavy chains per cluster, and the previously measured density of Myo2 myosin-II heavy chains (~3000 Myo2 heavy chains in a ~10 µm long ring, ref. 1).

(C) Value in protoplasts is estimated in ref. 2. Value in cell ghosts is chosen such that at t = 40 min 60% of F-actin in the ring is lost compared to t = 0, consistent with previous experiments[3].

(D) Value in protoplasts is estimated from FRAP measurements of Cdc12p[4]. In cell ghosts, actin loss in the presence of ATP and Phallacidin is negligible throughout constriction, suggesting formin molecules do not unbind the ring[3].

(E) Value in protoplasts is estimated from FRAP measurements of myosin light chain Cdc4p[5]. In cell ghosts, myosin loss is set to be consistent with Fig. S2a.

(F) Obtained from ref. 6.

(G) Estimated in ref. 7 using node motions measured there.

(H) Estimated from ref. 8. See ref. 9 for details.

(I) The coiled coil tail of Myo2 is 65 nm long[10]. The capture radius in the simulation is the tail length plus 15 nm to account for the length of the head and neck domain.

(J) Obtained from refs. 11, 12.

(K) Chosen so that the aggregation time observed in simulations and experiments in *S. japonicus* ghosts are similar.

(L) Estimated in ref. 2.
